# Genome-wide association study of flowering time reveals complex genetic heterogeneity and epistatic interactions in rice

**DOI:** 10.1101/2020.06.22.164616

**Authors:** Chang Liu, Yuan Tu, Shiyu Liao, Xiangkui Fu, Xingming Lian, Yuqing He, Weibo Xie, Gongwei Wang

**Affiliations:** National Key Laboratory of Crop Genetic Improvement and National Center of Plant Gene Research (Wuhan), Huazhong Agricultural University, Wuhan, China

**Keywords:** Flowering time, GWAS, epistatic interactions, genetic heterogeneity, rice

## Abstract

Since domestication, rice has cultivated in a wide range of latitudes with different day lengths. Selection of diverse natural variations in heading date and photoperiod sensitivity is critical for adaptation of rice to different geographical environments. To unravel the genetic architecture underlying natural variation of rice flowering time, we conducted a genome wide association study (GWAS) using several association analysis strategies with a diverse worldwide collection of 529 *O. sativa* accessions. Heading date was investigated in three environments under long-day or short-day conditions, and photosensitivity was evaluated. By dividing the whole association panel into subpopulations and performing GWAS with both linear mixed models and multi-locus mixed-models, we revealed hundreds of significant loci harboring novel candidate genes as well as most of the known flowering time genes. In total, 127 hotspots were detected in at least two GWAS. Universal genetic heterogeneity was found across subpopulations. We further detected abundant interactions between GWAS loci, especially in *indica*. Functional gene families were revealed from enrichment analysis of the 127 hotspots. The results demonstrated a rich of genetic interactions in rice flowering time genes and such epistatic interactions contributed to the large portions of missing heritability in GWAS. It suggests the increased complexity of genetic heterogeneity might discount the power of increasing the sample sizes in GWAS.

## Introduction

Cultivated rice (*Oryza sativa* L.), a major cereal especially in Asia, consists of two subspecies, *indica* and *japonica*. Since its domestication, rice has cultivated in a wide range of latitudes with different day lengths, from tropical to temperate regions. As a facultative short day (SD) plant, rice flowering can be promoted under SD conditions, whereas be repressed under long day (LD) conditions. A critical determinant for adaptation of rice to different geographical latitudes is the breeding selection of diverse natural variations in photoperiod sensitivity (Izawa 2007). For example, to ensure successful pollination and seed setting, rice cultivars grown in northern temperate regions need to flower very quickly under LD conditions during a short summer period, showing much less photoperiod sensitivity than those in other regions. A complex genetic network controlling rice flowering time in response to photoperiod has been elucidated. It shows similarities as well as divergences to that in *Arabidopsis*. In rice, *OsGI, Hd1* and *Hd3a* are determined as orthologs of *Arabidopsis GI, CO* and *FT*, respectively (Hayama et al. 2003; Kojima et al. 2002; Yano et al. 2000). Although these components are conserved, their regulation mode has been modified. In *Arabidopsis CO* only activates transcription of the florigen *FT* and promotes flowering under LD conditions. In rice, *OsGI* regulates the expression of *Hd1* and *OsMADS51. Hd1* has dual function, repressing *Hd3a* expression and flowering under LD while promoting flowering under inductive SD condition (Hayama et al. 2003). *Hd1* functions as a flowering repressor through interaction with Ghd7 (Nemoto et al. 2016). In contrast to *Arabidopsis*, rice can flower under non-inductive LD condition by inducing expression of a second florigen *RFT1* (Komiya et al. 2009). In addition, a unique *Ehd1* pathway, absent in *Arabidopsis*, was first discovered by Doi et al. (2004). *Ehd1* encodes a B-type response regulator and up-regulates florigen gene expression. It was shown that most LD repressors, such as *Ghd7, DTH8*/*Ghd8* and *OsMADS56*, act through *Ehd1* by reducing its expression and that of its downstream florigenc genes (Ryu et al. 2009; Wei et al. 2010; Xue et al. 2008; Yan et al. 2011). Under LD, Hd1 is a strong repressor of Ehd1 (Du et al. 2017; Goretti et al. 2017; Nemoto et al. 2016). *Ehd1* and *Ghd7* set critical day length for *Hd3a* florigen expression. *Ehd1* induction is mediated by *OsGI* and blue light at dawn, while *Ghd7* expression at dawn prevents *Ehd1* expression. However, under SD condition inducibility of *Ghd7* shifts to the dark phase and de-represses *Ehd1* (Itoh et al. 2010). Recently, several regulators which could promote *Ehd1* expression were also identified. *OsId1/RID1/Ehd2* acts as a master switch for flowering promotion under LD by up-regulating *Ehd1* expression (Matsubara et al. 2008; Park et al. 2008; Wu et al. 2008). *Ehd3* down-regulates *Ghd7* transcription, thus allowing *Ehd1* up-regulation under LD condition (Matsubara et al. 2011). *OsMADS50* strongly promotes *Ehd1* expression in LD condition, and *OsMADS50, DTH3* and *Hd9* are likely multiple alleles (Bian et al. 2011; Ryu et al. 2009). *Ehd4*, another activator of *Ehd1*, encodes a nucleus localized CCCH-type zinc finger protein unique to rice (Gao et al. 2013).

Although natural variation in flowering time has been studied extensively in rice, most of them was performed based on bi-parental quantitative trait locus (QTL) linkage mapping approach, with very limited range of allelic diversity and genomic resolution. In present study, using a diverse worldwide collection of 529 *O. sativa* accessions re-sequenced on the Illumina HiSeq 2000, the genetic architecture of natural variation in rice flowering time was characterized through GWAS with several association analysis strategies. Heading date and photosensitivity were investigated, and hundreds of significant association loci were identified. We detected 127 genomic hotspots associated with variation of rice flowering time by dividing the whole study population into subpopulations and using both linear mixed models and multi-locus mixed-models. We further analyzed interactions between loci which had been identified by GWAS in at least one (sub)population. A rich of genetic heterogeneity and epistatic interactions between flowering time genes were revealed in rice.

## Materials and Methods

### The association panel

The association panel consisted of a diverse collection of 529 *O. sativa* accessions including both core/ mini core collections and parental lines in breeding program. The details about sequencing, SNP identification, and imputation were described in Xie et al. (2015). The population structure of the whole association panel was inferred using ADMIXTURE (Alexander et al. 2009). The set of 529 rice accessions was classified into 98 *indica I* (*IndI*), 105 *indica II* (*IndII*), 92 *indica* intermediate, 91 temperate *japonica* (*TeJ*), 44 tropical *japonica* (*TrJ*), 21 *japonica* intermediate, 46 *Aus*, 14 intermediate group (*VI*), and 18 intermediate. Information about the accessions, including accession name, country of origin, longitude and latitude origin, and subpopulation identity, has been reported previously (Wang et al. 2015; Xie et al. 2015) and is available at the RiceVarMap (http://ricevarmap.ncpgr.cn).

### Field experiments and phenotyping

Field trials were carried out in three environments. The rice seeds were sown in the Experimental Station of Huazhong Agricultural University, Wuhan (central China, 30°28’N), in May of 2011 and 2012, and additionally in the Experimental Station of Lingshui County of Hainan Island (southern China, 18°48’N) in December of 2011. Seedlings about 25 days old were transplanted to the field. The field planting followed a randomized complete block design with two replications. Each plot consisted of four rows with 10 plants each. The planting density was 16.5 cm between plants in a row, and the rows were 26 cm apart. Field management, including irrigation, fertilizer application and pest control, followed essentially the normal agricultural practice.

Heading dates were recorded as the number of days from sowing to the time when the first panicles emerged above the flag leaf sheathes for half of the individuals in an accession. Heading dates in three environments for the 529 accessions were collected during the summer of 2011 and 2012 in Wuhan under a long-day condition (13.5∼14.2h) and during the spring of 2012 in Lingshui under a typical short-day condition (11.0∼12.5h). The correlation coefficient between two years of heading dates in Wuhan is 0.86, while the correlation coefficients between Hainan and Wuhan were 0.41 and 0.43, respectively.

### Genome-wide association analyses

Only SNPs with MAF ≥ 0.05 and the number of accessions with the minor allele ≥ 6 in a (sub)population were used to carry out GWAS. There are 2,046,642, 2,671,688, 2,767,191, 1,041,514, 1,857,866, and 3,916,415 SNPs used in GWAS for subpopulations of *IndI, IndII, Indica* (including *IndI, IndII*, and *indica* intermediate), *TeJ, Japonica* (includeing *TeJ, TrJ*, and *japonica* intermediate) and the whole population, respectively. Totally, 4,634,871 SNPs were involved in at least one GWAS. We performed GWAS using the linear mixed model (LMM) and the simple linear regression model (LR) provided by FaST-LMM program (Lippert et al. 2011) and multi-locus mixed-model (MLMM) from an R script provided by Segura et al. (2012). Population structure was modeled as a random effect in LMM using the kinship (K) matrix and we found that it was enough to control spurious associations, for the genomic inflation factor was near one in all GWAS. The evenly distributed random SNP set for analyzing population structure was used to calculate K. The kinship coefficients were defined as the proportion of identity genotype for the 188,165 randomly selected SNPs for each pair of individuals (Zhao et al. 2007). To reduce the computation burden, after obtaining association results of LR and LMM for each SNP, we split the genome into 50 kb regions and selected ten SNPs with the strongest association signals detected by LR and ten SNPs by LMM in the region. Ten SNPs in each 50 kb regions were usually included most of SNPs with *P*-value of LMM <0.01 in our GWASs. Association studies using MLMM were carried out using these SNPs. A uniform threshold *P*=5.0×10^−6^ was set to identify suggestive significant association signals by LMM (Chen et al. 2014; Wang et al. 2015). To obtain independent association signals, multiple SNPs exceeding the threshold in a 5 Mb region were clustered by *r*^*2*^ of LD ≥0.25 and SNPs with the minimum *P*-value in a cluster were considered as lead SNPs.

### Gene nomenclature

Rice genes with CCT domain and their nomenclature were obtained from Cockram et al. (2012). The nomenclature of MADS box genes was according to the annotation version 6.1 of genomic pseudomolecules of *japonica* cv. Nipponbare from Michigan State University (MSU).

### Epistatic interactions analysis

The analysis of two-locus interactions was carried out only between significant lead SNPs identified by LMM or MLMM. We fitted the following linear mixed model to identify two-locus interactions:

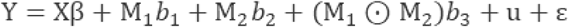

Where Y is a vector of a phenotype. The X is a matrix of fixed effects excluding SNPs, M_1_ and M_2_ are the selected two SNPs, 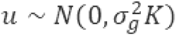 is a random effect, and ε is the noise term.

### Gene ontology (GO) enrichment analysis

The GO classifications of rice genes were downloaded from Gramene (http://www.gramene.org). Only the terms in the biological process category were used for GO analysis. The R package topGO was used to carry out GO enrichment analysis using Fisher’s exact test and the weight method (Alexa et al. 2006).

### Gene family enrichment analysis of GWAS hotspots

The Pfam domain information of rice genes from the MSU annotation version 6.1 was used in this analysis. Only gene-Pfam hits with E-value ≤ 1 × 10^−10^ were considered. A total of 472 Pfam domains owned by at least ten and at most 150 genes were used for enrichment analysis. A software INRICH was used to carry out interval-based enrichment analysis for GWAS hotspots (Lee et al. 2012).

## Results

### GWAS of heading date and photosensitivity in rice

Genome-wide association analyses were performed separately in the whole population and in the *IndI, IndII, indica* (consisting of *IndI, IndII* and *indica* intermediate), *TeJ*, and *japonica* (consisting of *TeJ, TrJ* and *japonica* intermediate) subpopulations for each environment. A total of 156 lead SNPs (the SNP with the lowest *P* value in a region) corresponding to 131 genomic clusters (adjacent lead SNPs in less than 100 kb were considered as a cluster) were detected in at least one population for the three datasets of heading date at linear mixed models (LMM). The analyses using multi-locus mixed-model (MLMM) based on the extended Bayesian information criterion (EBIC) detected a total of 234 lead SNPs with 198 genomic clusters. The details about these significant association signals are listed in Table S1 (for LMM) and Table S2 (for MLMM). The quantile-quantile plots and Manhattan plots for heading date of Wuhan_2012 in the whole population are illustrated in Fig. 1 as an example.

**Fig. 1.**
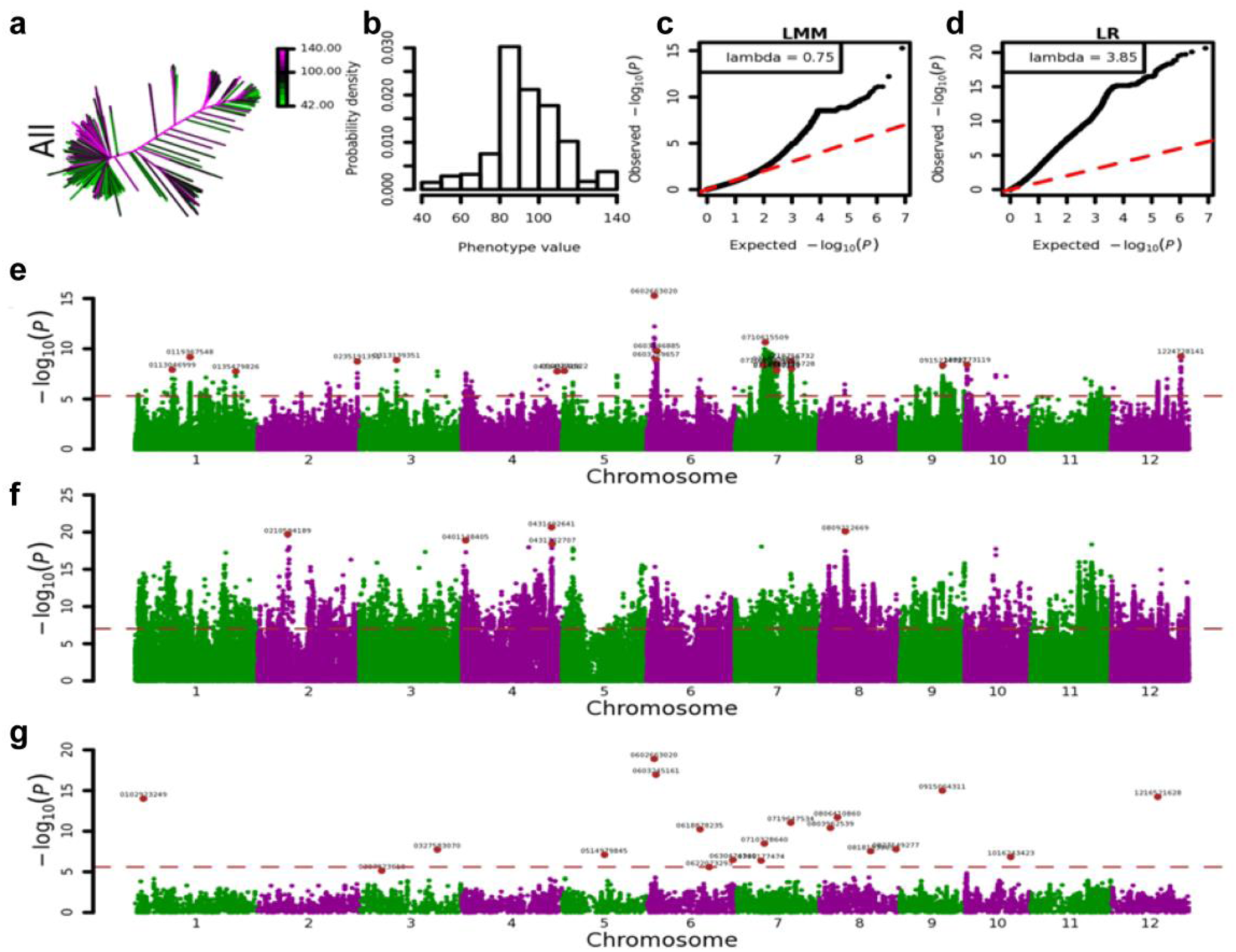
GWAS for heading date of Wuhan_2012 in the whole association population. (a-b) The heatmap (a) and histogram (b) distribution of heading date of Wuhan_2012 in 529 accessions. (c-d) Q-Q plot of the expected null distribution and the observed *P*-value using the linear mixed model (c) and the simple linear regression model (d). (e-g) Genome-wide *P*-values for the linear mixed model (e), simple linear regression model (f), and multi-locus mixed-model (g). The horizontal dashed line indicates the significance thresholds set as *P*=5.0×10^−6^ by LMM. The SNP positions of representative peak signals were denoted.

To search for candidate genes, the significant GWAS signals were compared with the positions of known or putative genes involved in flowering time pathway. We found that some known genes associated with rice flowering time, such as *Ghd7, Hd9* (*DTH3*), *Ehd1, OsMADS51*, and *SRT5*, were located within 50 kb adjacent to the identified lead SNPs (Table 1). Additional loci near known genes *Ghd8* and *Hd9* (*DTH3*), which were not significant at LMM model, could be detected using MLMM. *Hd9* (*DTH3*) was detected by both LMM and MLMM in short-day condition using the whole population or the *IndII* subpopulation, but only be detected by MLMM when using all *indica* accessions (Table 1). There were three independent lead SNPs for the three detections, which suggested allelic heterogeneity in *Hd9* (*DTH3*) in different subpopulations. Genes with CCT (CONSTANS, CO-like, and TOC1) or MADS box domains are usually associated with variations of rice flowering time. We found that three CCT genes (*OsPRR59, OsP* and *OsCMF10*) and two MADS genes (*OsMADS87* and *OsMADS30*) were also located close to the identified lead SNPs. These genes have not been reported yet but should be regarded as important candidates for flowering time in rice (Murakami et al. 2005; Tsuji et al. 2011). Lead SNPs near *OsMADS87* and *OsMADS30* passed the suggestive threshold in short-day condition in *indica* accessions by LMM, but they were only detected by MLMM in the whole population (Table 1), which suggests that LMM and MLMM might have complementary properties in GWAS. Other candidate genes for GWAS loci of heading date were also suggested in Tables S1 and S2.

**Table 1.**
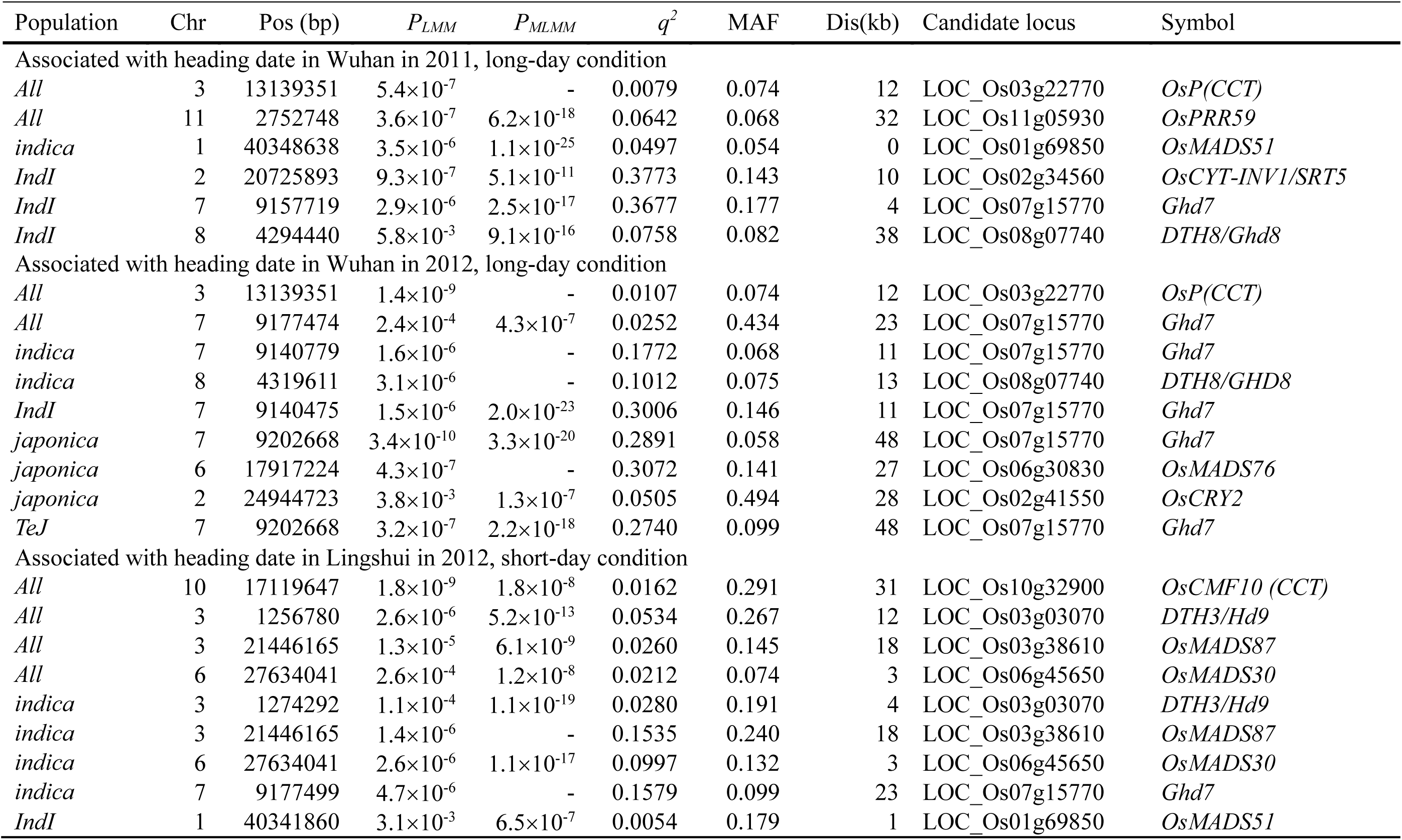
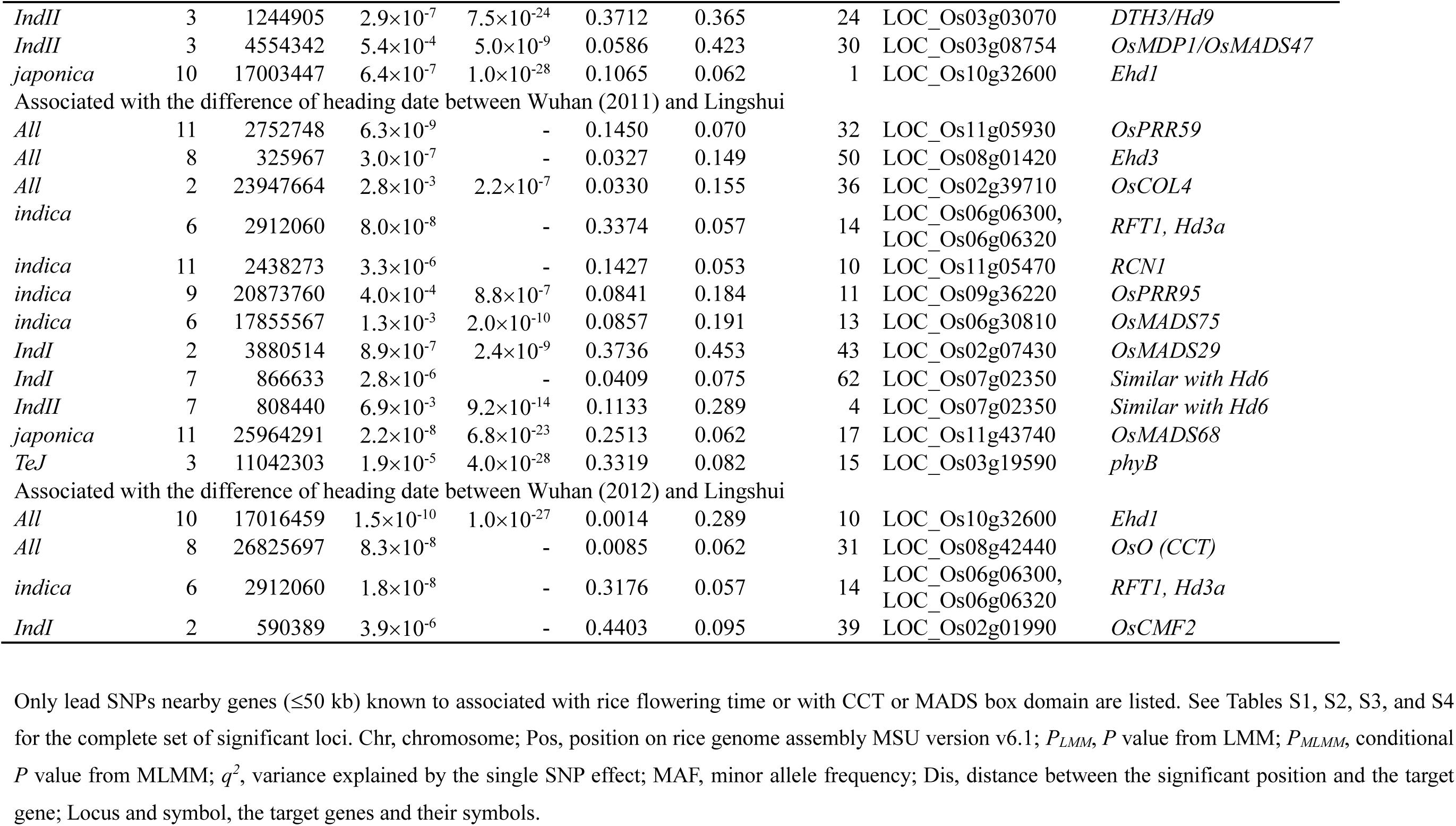
A subset of significant lead SNPs near known genes identified by GWAS for rice flowering time in Wuhan and Lingshui

Furthermore, the photosensitivity of accessions was investigated and used as a derived trait for GWAS. The differences of heading date between Wuhan and Lingshui were used to measure the photosensitivity. Totally 103 lead SNPs corresponding to 86 genomic clusters were detected by LMM (Table S3) and 25 of them were located near lead SNPs detected for heading date (<100kb). The analyses using MLMM detected a total of 184 lead SNPs with 168 genomic clusters (Table S4), 37 of which were located near lead SNPs detected for heading date (<100kb). Additional significant loci near known genes were detected for photosensitivity (Table 1), which suggests an important role of environment interactions in rice heading date. The CCT gene *OsPRR95* and three MADS box domain genes (*OsMADS75, OsMADS29* and *OsMADS68*) were important candidates for the photosensitivity in rice. Intriguingly, we did not detect association signal for *Hd6*, but signals near a gene (LOC_Os07g02350) similar to *Hd6* were detected by LMM in *IndI* and MLMM in *IndII* for difference of heading date between Wuhan and Lingshui in 2011 (Table 1). Compared with Hd6, LOC_Os07g02350 has 147 more amino acids at the N-terminal, and only 7 amino acids are different in the homologous region (Fig. S1). Other candidate genes for GWAS loci of photosensitivity were also suggested in Tables S3 and S4. The quantile-quantile plots and Manhattan plots for the differences of heading date between Wuhan_2012 and Lingshui in the whole population are illustrated in Fig. 2 as an example.

**Fig. 2.**
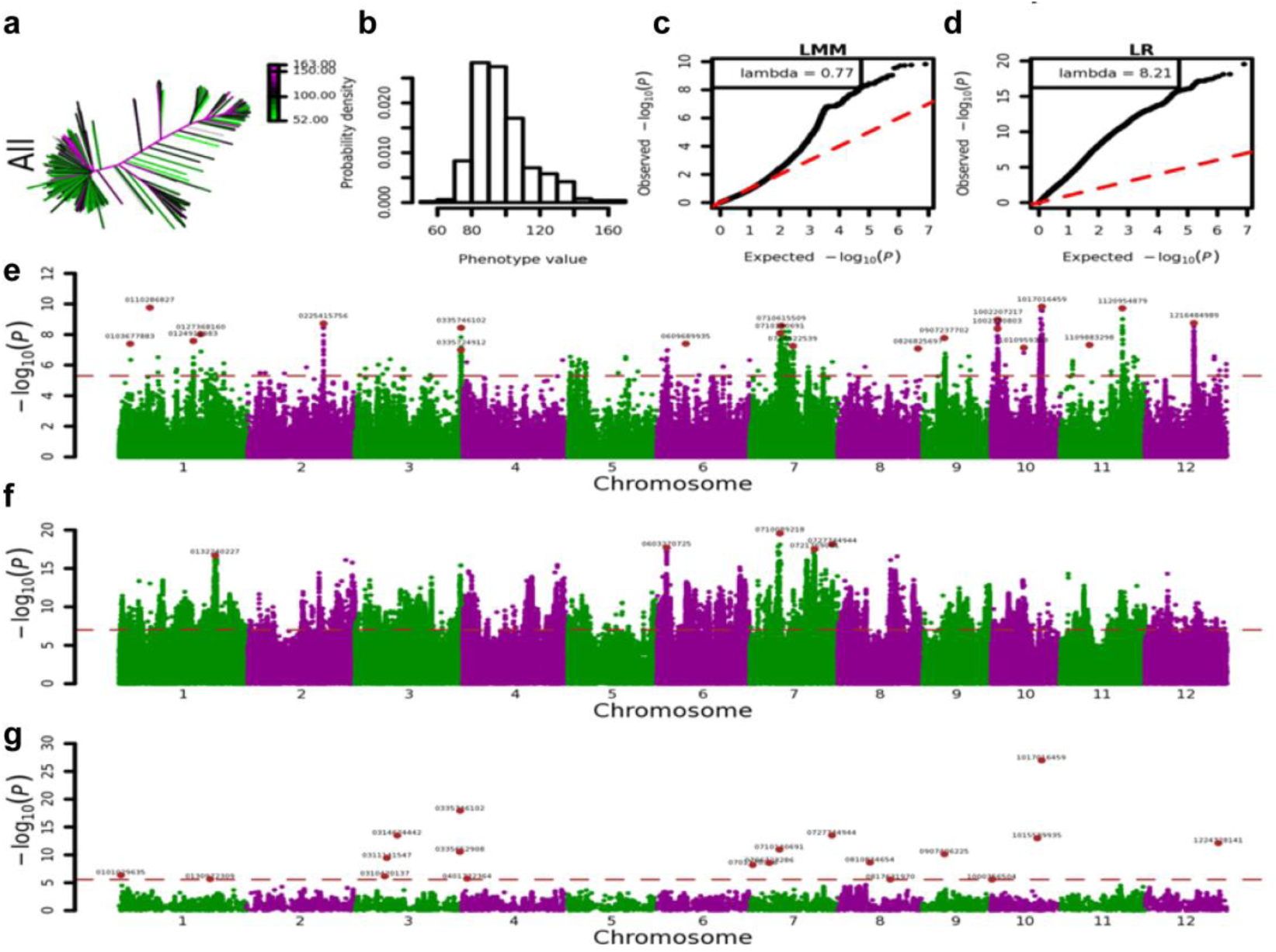
GWAS for the differences of heading date between Wuhan_2012 and Lingshui in the whole association population. (a-b) The heatmap (a) and histogram (b) distribution of the derived trait in 529 accessions. (c-d) Q-Q plot of the expected null distribution and the observed *P*-value using the linear mixed model (c) and the simple linear regression model (d). (e-g) Genome-wide *P*-values for the linear mixed model (e), simple linear regression model (f), and multi-locus mixed-model (g). The horizontal dashed line indicates the significance thresholds set as *P*=5.0×10^−6^ by LMM. The SNP positions of representative peak signals were denoted.

Besides known flowering time genes encoding proteins, we also observed that a lot of microRNAs were located near lead SNPs. miR156 (Xie et al. 2006), miR172 (Aukerman and Sakai 2003), miR159 (Achard et al. 2004), and miR399 (Kim et al. 2011), which have been shown to play important roles in flowering time in rice or *Arabidopsis*, were located near lead SNPs (Tables S1, S2, S3, and S4).

Taken together, there was a total of 248 lead SNPs detected by LMM (around ten lead SNPs were detected in average for each analysis) corresponding to 194 genomic clusters. Among them, thirty lead SNPs reoccurred in at least two GWASs. If taking different SNPs in a cluster as a common causal gene, there were 54 clusters detected in at least two GWASs, containing 108 lead SNPs. To estimate the genome-wide error rate in the GWASs, we performed 100 permutations for each analysis, with the details of the threshold shown in Table S5. The permutation results suggest that there were 2.37 false positives in one GWAS. The analyses using MLMM detected a total of 416 lead SNPs corresponding to 333 genomic clusters, of which 143 SNPs were near lead SNPs detected by LMM (<100kb). The selected optimal model of MLMM for each GWAS contained fourteen SNPs in average, ranging from three for some GWAS using *TeJ* to nineteen for the whole population. We observed that lead SNPs detected by LMM were enriched with lower MAF comparing with MLMM (Fig. S2). When merging the SNPs detected by MLM and MLMM together, there were a total of 611 SNPs forming 429 genomic clusters. Among them, 127 clusters were detected in at least two GWASs, and were thus referred to as hotspots. The 127 hotspots contained 309 lead SNPs, and 95 of them were detected by at least one LMM (Table S6). Because the less stringent threshold used might increase the risk of including false positives, we then focused on loci located in the 127 hotspots for further analysis.

### Universal genetic heterogeneity across subpopulations revealed by GWAS of rice flowering time

Most of the 127 hotspots contained multiple SNPs detected from different GWASs. We examined 151 SNPs located in the hotspots and detected by LMM in at least one GWAS, and found that only 44 SNPs showed polymorphic in all five subpopulations, indicating that a lot of loci were differentiated only in some of the subpopulations. Only 40 of the 151 SNPs exceeded the arbitrary threshold of *P*_*LMM*_ ≤ 5 × 10^−6^ for more than one subpopulation, and no SNP exceeded the threshold in both *indica* (*IndI, IndII* and *indica* intermediate) and *japonica* (*TeJ* and *japonica* intermediate) subpopulations simultaneously, suggesting the presence of universal genetic heterogeneity in rice flowering time. However, 102 hotspots were detected in more than one subpopulation, illustrating that there were multiple functional haplotypes in most of the hotspots. Interestingly, there were multiple independent lead SNPs, which were tightly close to each other and detected in one GWAS, in five clusters (Table S7). We found that such close lead SNPs detected in a GWAS were with opposite effects and resulted from independent variations (Table S7). Since the conditional *P* values for two SNPs decreased dramatically when they were both included in a model, they could not be detected by MLMM. Further researches were needed for distinguishing whether these were variations in different but tightly linked genes or independent variations of common genes.

### Abundant epistatic interactions revealed by GWAS of rice flowering time

Since the *indica* population for GWAS contains both *IndI* and *IndII* while *japonica* includes *TrJ* and *TeJ*, we examined if loci detected in a subpopulation could be also detected in the union populations. Interestingly, for lead SNPs located in the 127 hotspots, we observed that there were 27 SNPs with *P*_*LMM*_ ≤ 5 × 10^−5^ in *TeJ* and 24 of them were also with *P*_*LMM*_ ≤ 5 × 10^−5^ in *japonica*. In contrast, there were 27 and 30 SNPs with *P*_*LMM*_ ≤ 5 × 10^−6^ in *IndI* and *IndII*, but only two and six of them were with *P*_*LMM*_ ≤ 5 × 10^−5^ in *indica*, respectively. These observations indicated that the increased complexity of genetic background might neutralize the effect through increasing the number of samples in GWAS.

For example, we found that *Ghd7*, a gene suppresses flowering under long-day conditions by suppressing the expression of *Ehd1*, was detected in *IndI* (Fig. 3a) but not in *indica* (Fig. 3b) in Wuhan_2011. Since the functional variation of *Ghd7* in *indica* is mainly a complete deletion of GHD7 (Xue et al. 2008), all lead SNPs should be resulted from indirect associations with the deletion. We examined the lead SNP sf079137593 near *Ghd7* detected by LMM in *IndI* in Wuhan_2011, and found that the SNP could only represent the deletion variation of GHD7 in *IndI* but not in other *indica* accessions (*r*^*2*^= 0.70 and 0.31 in *IndI* and *indica*, respectively. See Table S8 for details), suggesting the gains of GWAS in subpopulations. However, another SNP sf079157719, which was detected by MLMM and represent the deletion of GHD7 well in *indica* (*r*^*2*^= 0.93 and 0.77 in *IndI* and *indica*, respectively), still couldn’t be significantly detected in GWAS of *indica* either.

**Fig. 3.**
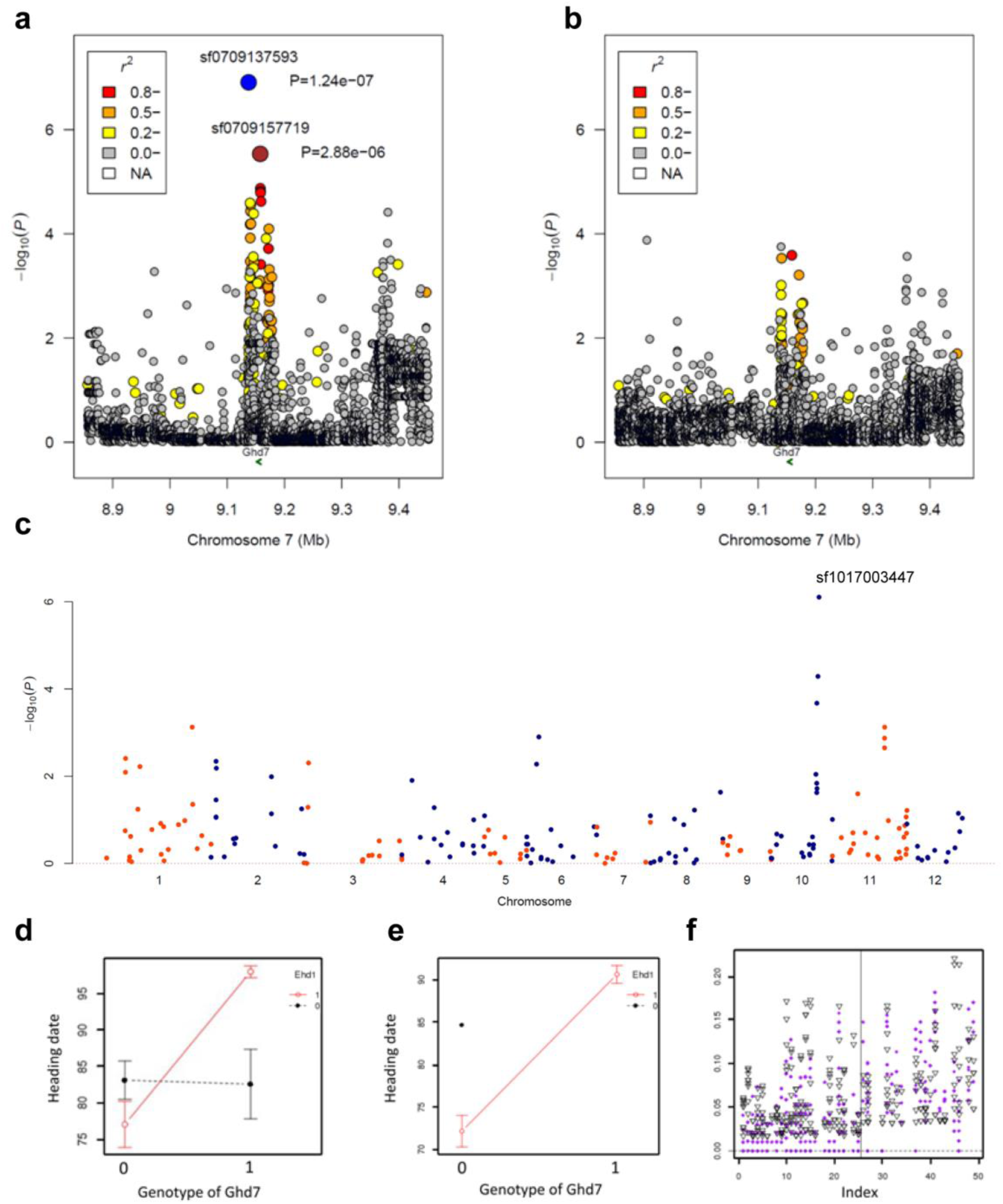
The GWAS results near *Ghd7* region for rice flowering time and epistatic interaction between *Ghd7* and *Ehd1*. (a-b) Association results near *Ghd7* for heading date in Wuhan_2011 in *IndI* (a) and *indica* (b). The blue point denotes the lead SNP sf079137593. The colors of the other points represent the linkage disequilibrium for the lead SNP. The arrow represents the position of *Ghd7*. (c) The result of scanning the interactions of *Ghd7* towards other lead SNPs. (d-e) epistatic interaction between *Ghd7* and *Ehd1*. A lead SNP sf1017003447 near *Ehd1* had the most significant epistatic interaction with *Ghd7* in *indica* (d). In contrast, sf1017003447 in *IndI* was nearly fixed (e). The distribution of the minor allele frequencies for the interacted SNPs. Each column represents the ten most significant SNPs interacted with a certain SNP. Before the vertical line: twenty-five SNPs with PLMM ≤ 5.0 × 10^−6^ in *IndI* and PLMM ≤ 5.0 × 10^−5^ in *indica*. The inverted triangles represent the minor allele frequencies of SNPs showing significant interactions in *indica*, while the points in purple is the minor allele frequencies of these interacted SNPs in *IndI* (obviously lower than in *indica*). After the vertical line: twenty-four SNPs with PLMM ≤ 5.0 × 10^−5^ in both *TeJ* and japonica. The inverted triangles represent the minor allele frequencies of SNPs showing significant interactions in japonica, while the points in purple is the minor allele frequencies of these interacted SNPs in *TeJ*. (f).

As many epistasis between loci controlling heading date in rice have been discovered, we tried to examine whether such epistatic interactions could be detected in our data and might lead to missing of detecting *Ghd7* in *indica*. To reduce computational requirements and false positives, only the 611 lead SNPs detected by LMM or MLMM methods in our GWASs were used to analyze interactions. The interactions of any two loci were examined in *indica* population if the minimum numbers of individuals of all four genotype combinations (denotes as *Nc*) were no less than five. When scanning the interactions of *Ghd7* towards other lead SNPs, we found that a lead SNP sf1017003447 near *Ehd1* had the most significant *P* value of epistatic interaction (Fig. 3c), which was consistent with previous genetic researches (Xue et al. 2008). When only considering single locus, both lead SNPs were not significant (*P*=0.084 for sf079157719 and *P*=0.236 for sf1017003447). But when considering interaction, not only the interaction itself was extremely significant (*P* = 2.13 × 10^−4^) but also the two loci were significant (*P*=0.027 for *Ghd7* and *P*=0.029 for *Ehd1*) (Fig. 3d). We also investigated the situation in *IndI*, and found that the lead SNP sf1017003447 in *IndI* was nearly fixed (Fig. 3e), therefore, the effect of interaction did not cause trouble for the identification of *Ghd7* in *IndI*.

We further inspected the ten most significantly interacted SNPs in *japonica* for each of the 24 SNPs with *P*_*LMM*_ ≤ 5 × 10^−5^ in both *TeJ* and *japonica*. These associations are likely not affected by population differentiation. Indeed, there were 112 interactions with *P* ≤ 0.01 in *japonica*. Ninety of them (80.4%) were with *Nc* ≤ 5 in *TeJ* and 31 (27.7%) were also significant with *P* ≤ 0.01 in *TeJ*. However, for the 25 SNPs with *P*_*LMM*_ ≤ 5 × 10^−6^ in *IndI* and *P*_*LMM*_ ≤ 5 × 10^−5^ in *indica* (i.e., detected in *IndI* only, These associations are likely affected by population differentiation), there were 191 interactions with *P* ≤ 0.01 in *indica* but only 36 of them (18.9%) were with *Nc* ≤ 5 in *IndI* and none of them was significant with *P* ≤ 0.01 in *IndI*. The distribution of the frequencies of *Nc* for these SNPs was shown in Fig. 3f, which indicated that for interactions significant in *indica*, a lot of genotype combinations did not exist in *IndI*. Similar situations were also observed in *IndII*. But for interactions significant in *japonica*, many of them were also found in *TeJ*. We thus propose that the new emerging interactions in *indica* might account for the missing of the significant lead SNPs that were able to be detected in *IndI* or *IndII*. Such results also suggest that epistatic interaction is an important part of GWAS and should be considered as the large component of missing heritability in GWAS (Brachi et al. 2011).

### Functional gene families revealed from enrichment analysis of GWAS hotspots for rice flowering time

Based on the 127 GWAS hotspots, we assessed whether genes with certain conservative domains enriched in genomic regions near lead SNPs using INRICH (Lee et al. 2012). We found that CCAAT binding factors (PF00808), MADS-box transcription factors (PF00319), Myb-like DNA-binding domain proteins (PF00249), and CCT domain proteins (PF06203), which are usually harbored by known rice flowering time genes, were ranked in the top of enrichment analysis with *P* ≤ 0.1 (Table 2). The top significant enriched Pfam protein domains contained a lot of transcription factors, suggesting that transcript regulation might be the main way to regulate flowering in rice. Intriguingly, we found that besides *Ghd8* which encodes a putative HAP3 subunit of the trimeric HAP2/HAP3/HAP5 complex (also known as CCAAT binding factor), three genes (LOC_Os02g07450, LOC_Os03g63530, and LOC_Os06g45640) encoding HAP5 subunit were also located near lead SNPs. It is proposed that CCT domain proteins share similarity to HAP2 and might replace that component to form the trimeric complex (Laloum et al. 2012; Wenkel et al. 2006). It seems that our results provided evidences for this supposal, for all three components were significantly enriched in GWAS results. Genes encoding histones (PF00125) were enriched, which might indicate the important roles of histone modifications for regulating flowering time in rice (He et al. 2003). Genes encoding kinesin (PF00225) were also enriched. A recent study showed that a kinesin OsGDD1 acted as a transcription factor for the synthesis of the phytohormone gibberellin in rice (Li et al. 2011), suggesting a possible role of kinesin genes for regulating flowering time in rice. Further experiments were needed to elucidate whether the genes encoding the enriched Pfam domain were true positive.

**Table 2.** The top fifteen enriched Pfam protein domains detected from GWAS hotspots for rice flowering time

| Pfam domain | Total gene | No. in hotspot | *P* value | Description |
| --- | --- | --- | --- | --- |
| PF00808 | 24 | 4 | 0.017 | Histone-like transcription factor (CBF/NF-Y) |
| PF02364 | 14 | 3 | 0.017 | Glucan_synthase |
| PF01398 | 15 | 3 | 0.023 | JAB1/Mov34/MPN/PAD-1 ubiquitin protease |
| PF00702 | 55 | 6 | 0.026 | haloacid dehalogenase-like hydrolase |
| PF00319 | 55 | 6 | 0.029 | SRF-type transcription factor (DNA-binding and dimerisation domain) |
| PF01167 | 15 | 3 | 0.030 | Tubby protein |
| PF00225 | 48 | 6 | 0.033 | Kinesin motor domain |
| PF02170 | 21 | 3 | 0.041 | PAZ domain |
| PF00125 | 48 | 5 | 0.045 | Core histone H2A/H2B/H3/H4 |
| PF00249 | 146 | 11 | 0.049 | Myb_DNA-binding |
| PF00501 | 45 | 5 | 0.057 | AMP-binding |
| PF07762 | 31 | 3 | 0.063 | DUF1618 |
| PF00118 | 20 | 3 | 0.066 | TCP-1/cpn60 chaperonin family |
| PF06203 | 39 | 4 | 0.077 | CCT |
| PF00010 | 50 | 5 | 0.086 | Helix-loop-helix DNA-binding domain |

## Discussion

Heading date is a very important and complex trait that controls adaptation of rice varieties to their local environment and yield performance, and flowering time loci are often the targets of both natural and artificial selection. The trait is strongly affected by population structure, and the loci often exhibit complex forms of allele sharing and admixture in diverse germplasm. Previous GWAS studies in rice yielded only a few known loci associated with heading date, with some or even a large portion of known functional genes failed to be identified (Huang et al. 2012; Zhao et al. 2011). In present study, to unravel the genetic architecture underlying natural variation of rice flowering time, using a diverse worldwide collection of 529 *O. sativa* accessions as the GWAS platform, we adopted several association analysis strategies, which enabled us to detect hundreds of significant association loci. Heading date in three environments was recorded during the summer seasons of 2011 and 2012 in Wuhan (central China, 30°28’N) under a long-day condition (13.5∼14.2h) and during the spring season of 2012 in Lingshui (southern China, 18°48’N) under a typical short-day condition (11.0∼12.5h). Photosensitivity of rice accessions was evaluated (differences of heading date between long-day condition in Wuhan and short-day condition in Lingshui) and used as a derived trait for GWAS. Previous studies discovered that additional association signals can be detected when GWASs are performed on subpopulations and suggested the existence of genetic heterogeneity across subpopulations in rice (Huang et al. 2010; Huang et al. 2012; Zhao et al. 2011). In addition to the whole population, we also divided it into several subpopulations and performed GWAS. The whole genome scanning of all SNPs with MAF ≥ 0.05 in the target population was first carried out using both a simple linear regression (LR) and a linear mixed model (LMM) (Lippert et al. 2011). A recent study showed that the multi-locus mixed-model (MLMM) outperforms the single-locus model (Segura et al. 2012). We also performed stepwise mixed-model regressions to construct multi-locus models based on preliminary LMM and LR results. Our results suggest that LMM and MLMM might have complementary properties in GWAS. We revealed hundreds of significant loci harboring novel candidate genes as well as most of the known flowering time genes. According to literature, there are eight rice flowering time genes identified by map-based cloning, and five of them (*DTH3, Hd3a, Ghd7, Ghd8*, and *Ehd1*) were detected in at least two GWASs in our study. Although GWAS is a powerful tool, due to the limitation of population size, GWAS will show its limitations, which may lead to the occurrence of false-positive. How to reduce the occurrence of false-positive and improve the accuracy of detection remains to be further studied.

Genetic heterogeneity across subpopulations for flowering time was analyzed in depth in this study. We found that a lot of significant loci were differentiated only in some of the subpopulations. No lead SNPs exceeded the threshold in both *indica* (*IndI, IndII*, and *indica* intermediate) and *japonica* (*TeJ, TrJ*, and *japonica* intermediate) subpopulations simultaneously. Some close lead SNPs detected in a GWAS were even with opposite effects and resulted from independent variations. These results suggest the presence of universal genetic heterogeneity in rice flowering time. Further, it is interesting to note that, especially in *indica*, the increased complexity of genetic background might neutralize the effect through increasing the number of samples in GWAS. For epistatic interactions between flowering time loci significant in *indica*, a lot of genotype combinations did not exist in *IndI*, either in *IndII*. But for those interactions significant in *japonica*, many of them were also found in *TeJ*. We propose that the new emerging interactions in *indica* might account for the missing of the significant lead SNPs that were able to be detected in *IndI* or *IndII*.

In present study, besides known genes and microRNAs in flowering time regulation located around lead SNPs, a lot of candidate genes were also suggested for the GWAS loci. We found that CCAAT binding factors, MADS-box transcription factors, Myb-like DNA-binding domain proteins, and CCT domain proteins, which are usually harbored by known rice flowering time genes, was ranked in the top of enrichment analysis with *P* ≤ 0.1 in the 127 GWAS hotspots. Many genes with known domains associated with flowering time were first reported in our study and should be good candidates for further studies. We noticed that *Hd1*, the first map-based cloned flowering time gene in rice, which was observed to be corresponding to a ‘mountain range’ in GWAS by Zhao et al (2011), could not be even detected in our study. Instead, we observed a ‘mountain range’ in *Ehd1* region in several GWASs. This might be due to different rice lines and environments in different studies. For example, *Ghd7* was cloned by our laboratory and could be detected clearly in different seasons and populations but did not display signal in Zhao et al (2011). Such observations suggest calls for collaborations that a common set of rice lines should be planted in different places around the world and such collaborations would lead to detection of more functional genes.

## Authors’ contributions

WX and GW performed data analysis and wrote the manuscript; CL performed the final data analysis; YT, SL, XF, XL, and YH conducted field trials and collected the phenotypic data.

