## Supplementary Figures for "Genome-wide association study of flowering time reveals complex genetic heterogeneity and epistatic interactions in rice"

**Supplementary Data**

**Table S1** The details of the significantly associated SNPs for rice heading date detected by LMM.

**Table S2** The details of the selected SNPs for rice heading date detected by MLMM based on the extended Bayesian information criterion.

**Table S3** The details of the significantly associated SNPs for rice photosensitivity detected by LMM.

**Table S4** The details of the selected SNPs for rice photosensitivity detected by MLMM based on the extended Bayesian information criterion.

**Table S5** Significant threshold and genome-wide error rice (GWER) of GWAS for rice flowering time obtained from 100 permutations.

**Table S6** List of the 127 hotspots.

**Table S7** List of the five clusters regions with multiple independent lead SNPs, which were tightly close to each other and detected in one GWAS.

**Table S8** The genotype dataset of *Ghd7* region in *indica* accessions.

**Fig. S1** Homologous sequence alignment of Hd6 (LOC_Os03g55389) and Hd6-like (LOC_Os07g02350).

**Fig. S2** Lead SNPs detected by LMM were enriched with lower MAF compared with MLMM.

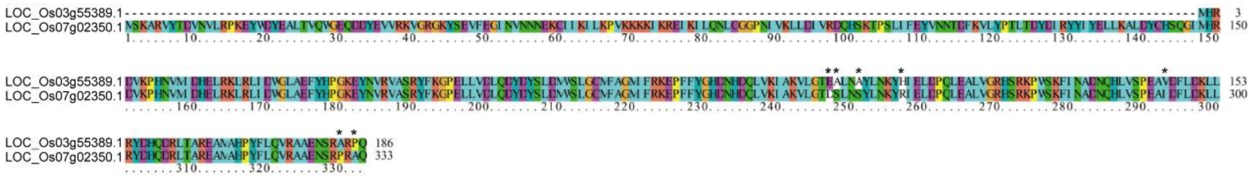


**Fig. S1** Homologous sequence alignment of Hd6 (LOC_Os03g55389) and Hd6-like (LOC_Os07g02350). * indicates the different amino acids in the homologous region.

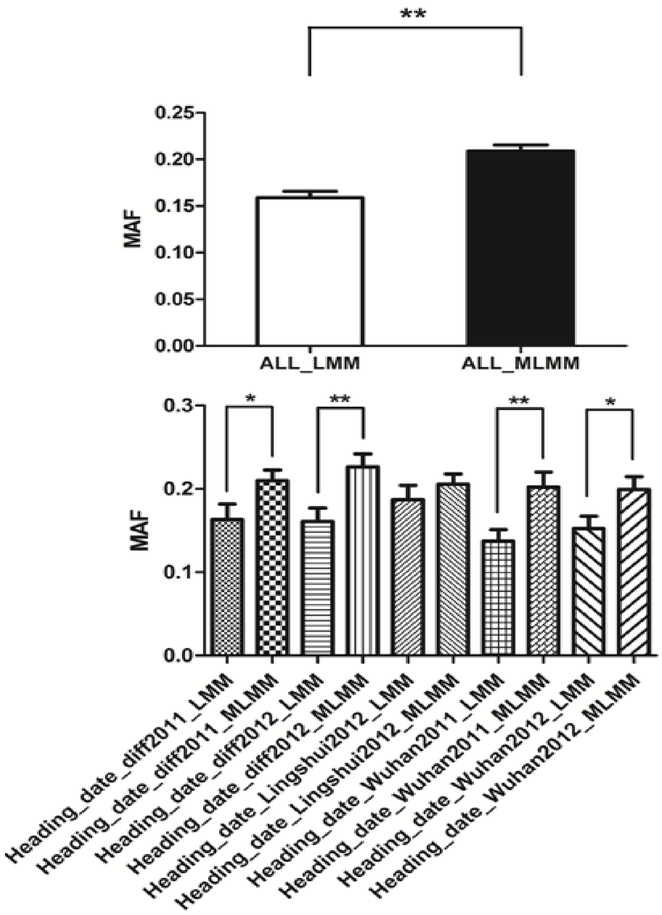


**Fig. S2** Lead SNPs detected by LMM were enriched with lower MAF compared with MLMM. MAF, minor allele frequency. ALL, the average of five sets of comparisons. * or ** indicates the differences of MAF between LMM and MLMM are significant at $P < 0.05$ or $P < 0.01$, respectively.
